# Translating insights from the seed metabolome into improved prediction for healthful compounds in oat (*Avena sativa L.*)

**DOI:** 10.1101/2020.07.06.190512

**Authors:** Malachy T. Campbell, Haixiao Hu, Trevor H. Yeats, Melanie Caffe-Treml, Lucía Gutiérrez, Kevin P. Smith, Mark E. Sorrells, Michael A. Gore, Jean-Luc Jannink

## Abstract

Oat (*Avena sativa* L.) seed is a rich resource of beneficial lipids, soluble fiber, protein, and antioxidants, and is considered a healthful food for humans. Despite these characteristics, little is known regarding the genetic controllers of variation for these compounds in oat seed. We sought to characterize natural variation in the mature seed metabolome using untargeted metabolomics on 367 diverse lines and leverage this information to improve prediction for seed quality traits. We used a latent factor approach to define unobserved variables that may drive covariance among metabolites. One hundred latent factors were identified, of which 21% were enriched for compounds associated with lipid metabolism. Through a combination of whole-genome regression and association mapping, we show that latent factors that generate covariance for many metabolites tend to have a complex genetic architecture. Nonetheless, we recovered significant associations for 23% of the latent factors. These associations were used to inform a multi-kernel genomic prediction model, which was used to predict seed lipid and protein traits in two independent studies. Predictions for eight of the 12 traits were significantly improved compared to genomic best linear unbiased prediction when this prediction model was informed using associations from lipid-enriched factors. This study provides new insights into variation in the oat seed metabolome and provides genomic resources for breeders to improve selection for health-promoting seed quality traits. More broadly, we outline an approach to distill high-dimensional ‘omics’ data to a set of biologically-meaningful variables and translate inferences on these data into improved breeding decisions.

## Introduction

The oat seed contains a diverse array of compounds that are beneficial for human health and nutrition (Gulvady et al. 2013). It is widely considered a healthy food due to its high soluble fiber content, which is unique among major cereals and has been shown to improve cardiovascular health and blood glucose levels (Gulvady et al. 2013; Kale et al. 2013). Oat is also a good source of protein (12.4-24.5% of seed weight), oil (3-11%), and a rich source of vitamins and minerals (Frey and Holland 1999; Gulvady et al. 2013). The oils found in the oat seed are primarily triglycerides, with palmitic, oleic, and linoleic acids being the primary fatty acids (Youngs 1978). In addition to the benefits from direct consumption, colloidal oatmeal and oat extracts have been used extensively as an effective topical medicine to treat skin dermatitis and reduce inflammation (Cerio et al. 2010; Kurtz and Wallo 2007). These benefits have been attributed to avenanthramides, flavonoids, tocopherol, polysaccharides, and lipids. Thus, the oat seed is a rich source of diverse compounds that have multifaceted effects on human health. To improve specific biochemical properties of oat, breeders must be provided with a suite of tools that allow these compounds to be quantified accurately at low cost, and genomic resources that improve selection for specific seed qualities.

Advances in biochemistry have provided a breadth of tools to query the metabolome and quantify known and unknown compounds (Dunn and Ellis 2005). Untargeted metabolomics can quantify 100s-1000s of metabolites in a sample, thus health-promoting and quality-related metabolites, and their intermediate or related compounds can be assessed with relative ease (Dunn et al. 2013; Christ et al. 2018). These data can be used to address basic biological questions regarding biochemical pathways that are represented in the data, and assess natural variation for these pathways. The effectiveness of these methods to characterize natural variation in the metabolome has been highlighted by several studies (Caspi et al. 2014; Chan et al. 2010; Matsuda et al. 2015; Slenter et al. 2018; Wu et al. 2018). Moreover, these data have been used as predictors, often alongside genomic data, to improve prediction for complex traits (Riedelsheimer et al. 2012; Guo et al. 2016; Xu et al. 2016).

Parsing these data to understand the biology of the seed metabolome can be challenging. Numerous databases are available that describe primary and secondary metabolic pathways, and are curated using information both across and within species (Kanehisa and others 2002; Wishart et al. 2020). Metabolites can be mapped to these pathways to determine which pathways and their products are enriched in a given set of samples. While these approaches provide greater confidence over unsupervised, data-driven approaches, in many cases only a fraction of the compounds quantified via untargeted metabolomics can be mapped to these pathways (Schrimpe-Rutledge et al. 2016; Cui et al. 2018). Unsupervised, data-driven approaches provide an attractive alternative that utilizes the data more completely. These approaches include co-expression-based analyses and factor analytic models among others. While coexpression-based analyses have been used extensively to characterize high-dimensional ‘omics’ data, these often require users to select several parameters that influence outcomes and may limit reproducibility (DiLeo et al. 2011; Langfelder and Horvath 2008). Factor analytic models, on the other hand, use a linear model to identify groups of strongly correlated metabolites. The underlying rationale for these approaches is that covariance among metabolites is driven by some unobserved (i.e., latent) underlying variable(s). With this approach, the matrix of metabolites is decomposed into a lower-dimensional linear combination of factor loadings, which describe how each latent factor contributes to each compound, and a set of factor scores that ascribe a phenotypic value for all individuals for a given latent factor. Thus, these frameworks have advantages from both biological and statistical perspectives. While in some respects factor analytic models achieve the same goal as others, such as principal component analysis (PCA) — providing a reduced rank representation of the data — the defining feature of factor analytic models is that latent factors are constructed to preserve correlation among groups of related metabolites. In PCA, new constructs are defined that preserve variance in the observed variables. Thus, constructs from factor analytic models can provide insight into biological processes driving covariation between phenotypes. Moreover, the lower-dimensional set of factor-scores can be treated as any other phenotype and reduce the multiple testing burden often associated with high-dimensional ‘omics’ datasets. With these frameworks we can address: (1) *What pathways are represented in the metabolome?* and (2) *How do these pathways and their products co-vary within a genetic population?*

Improving health or quality-related compounds requires decomposing phenotypic variation within the metabolome into genetic and non-genetic components, and utilizing these outcomes to inform selection decisions for quality-related phenotypes. Conventional linkage analysis or association mapping approaches have proven to be powerful approaches to identify genetic variants associated with variation in the metabolome (Chan et al. 2010; Eckert et al. 2012; Matsuda et al. 2015; Rowe et al. 2008; Wen et al. 2014; Xu et al. 2017). However, a much greater challenge is to translate genetic signal for health-promoting compounds, and related metabolites, to improve prediction and selection of new crop germplasm.

A number of studies have extended the conventional frameworks used for genomic prediction to accommodate prior biological information regarding genetic marker effects (Edwards et al. 2016; MacLeod et al. 2016; Speed and Balding 2014; Turner-Hissong et al. 2019). Although these approaches differ in how these data are treated, the motivation is similar for all —effects for variants that are more likely to be causative should be drawn from a different distribution than those lacking evidence for causality. Thus, prediction should be improved when effect sizes differ between genetic marker classes. The approaches described by Speed and Balding (2014) and Edwards et al. (2016) are essentially an extension of the genomic best linear unbiased prediction (gBLUP) framework, in which genomic markers are partitioned and are used to construct separate genomic relationship matrices for each random genetic effect. Genomic values for each individual are sampled from each distribution. The framework described by MacLeod et al. (2016) extends the Bayesian prediction framework, BayesR, and uses biological information to partition markers into classes (Erbe et al. 2012). Marker effects, rather than genomic values, are sampled from each distribution. In the context of the current study, if we know what metabolites are related to quality traits and have identified variants associated with these metabolites, genomic markers can be partitioned to define biologically informed marker-sets. These biologically informed marker-sets should be enriched for causal loci, and should improve prediction of genomic values.

In this study, we characterized the seed metabolomes of 375 oat lines and sought to identify loci that potentially influence (co)variation among many metabolites. Specifically, we sought to answer: (1) *What pathways or metabolite classes are enriched in the seed metabolome?* (2) *What are the genetic controllers of the metabolome?* and (3) *Can these data be leveraged to improve genomic prediction for seed quality traits?* To this end, we assayed the seed metabolome using untargeted LC-MS and GC-MS and used the empirical factor analysis approach described by Wang and Stephens (2018) to identify latent factors that generate covariance among many metabolites. GWAS was performed using this reduced set of latent phenotypes, and these outcomes were used to inform a multi-kernel genomic prediction model to predict seed quality traits in two independent studies. In summary, we extract meaningful basic biological insights from ‘omics’ data with limited annotations, and translate these outcomes to improve prediction for agriculturally important traits.

## Results

### Metabolite differences across subpopulations are primarily generated by drift

To characterize the metabolome of mature oat seed, we generated untargeted metabolite data using two mass spectroscopy (MS) pipelines (gas chromotography MS, GC-MS and liquid chromotography MS, LC-MS) for 367 diverse accessions (Supplemental File S1). The diversity panel consisted of 367 accessions that could be partitioned into six distinct genetic clusters using a *k*-means clustering approach (Fig. 1A and B; Fig. S1). Despite six clusters being identified, the degree of stratification within the population was minor. For instance, the first and second principal axes explained only 7.3% and 5.9% of the variance in genetic relationships, respectively (Fig. 1A and B). We quantified 1,668 metabolites (601 for GC-MS and 1,067 for LC-MS) across the 367 accessions. PCA of the metabolome dataset did not reveal any apparent clustering among accessions, and evidence of stratification between genetically-defined clusters was not visually apparent (Fig. 1C and D).

**Figure 1.**
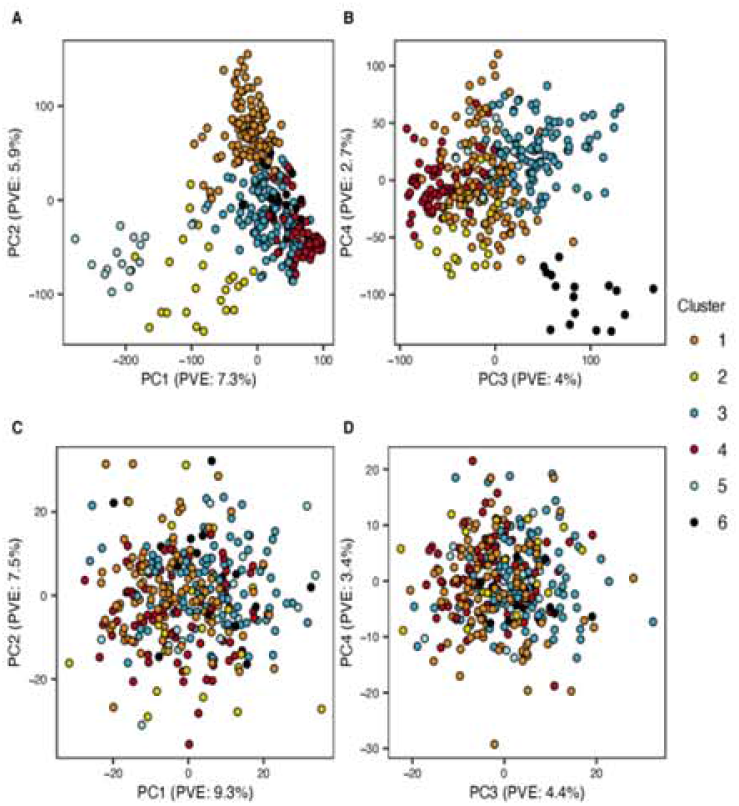
Principal component analysis of genotypic and metabolomic data. The first four principal components (PCs) of gentoypic data are shown in panels A and B, while the first four PCs of the metabolomic data are shown in panels C and D. Subpopulations that were defined based on *k*-means clustering of SNP marker data are indicated by different colored points. PC: principal component; PVE: percent variance explained

To determine whether individual metabolites differed among clusters, we performed a one-way ANOVA for each of the 1,668 metabolites (Supplemental File S2). Despite no strong differentiation of the metabolome between the six clusters, 41% of the 1,668 metabolites showed significant differences between one or more of the subpopulations (Benjamini-Hochberg adjusted *p*-value; *p*_*BH*_ < 0.01). We elucidated whether these differences were due to selection or drift by examining *P*_*ST*_, a measure of phenotypic divergence between populations, and compared these values to the distribution of genetic divergence (i.e., *F*_*st*_) for all loci (Storz 2002; Leinonen et al. 2013). This analysis revealed only 12 compounds with *P*_*st*_ values that were greater than 80% of the *F*_*ST*_ values, indicating that the majority of compounds differing between subpopulations diverged due to drift or weak selection. Only four of these compounds have annotations and were described as a putative steroidal glycosides, terpene glycoside, triterpenoid, and 1-benzopyran. These results suggest that the divergent metabolites are largely due to drift rather than selection.

### Latent factor model selection

Given that only a fraction of the metabolites quantified in our population were annotated, we leveraged the correlation between annotated and unannotated metabolites to infer biological processes in the oat seed with the rationale that compounds participating in a related biological process will be correlated. We used an unsupervised learning approach that distills the covariance among a set of observed variables into a lower dimensional set of unobserved constructs that may cause this covariance. In a biological sense, these latent factors may provide insights into the major biochemical features of the metabolome, and can be used to elucidate the genetic factors that shape the metabolome.

The covariance among the 1,668 metabolites was decomposed into a set of latent factors using the empirical Bayes matrix factorization (EBMF) approach described by Wang and Stephens (2018). This method constructs latent factors, which are defined by a linear combination of factor loadings and factor scores, by approximating the posterior sampling distribution for these parameters from the data (Wang and Stephens 2018). Three latent factor models that differed in the family of prior distributions (Laplace, point normal, and adaptive shrinkage) for factor loading and scores were evaluated, and the best model was selected based on the goodness-of-fit and predictive ability (Owen et al. 2016) (Table I, Fig. S2). The Laplace family of densities exhibited the lowest RMSE (0.970) and highest correlation between predicted and observed data (*r* = 0.520). The common covariance in the oat seed metabolome could be captured using 100 latent factors that collectively explained 58.82% of the total variance in the metabolite data.

**Table I.** Empirical Bayes matrix factorization model selection. Each model was fit using degressed BLUPs for 1,668 metabolites. Ad. Shr.: adaptive shrinkage family of densities described by Stephens (2016). Cross-validation (CV) was based on a 3-fold orthogonal CV described by (2018) and Owen et al.(2016) with ten independent resamplings. Point Nor.: point-normal family of densities which are a normal distribution with a point mass at zero; LL indicates log-likelihood; PVE: percent variance explained; 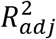: adjusted *R*^2^; 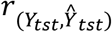 is the Pearson’s correlation between predicted and observed values for observations in the testing set; RMSE: root mean square error.

| EBNM Appr. | No. Fact. | LL | PVE | $R_{adj}^2$ | $r_{(Y_{tst}, \hat{Y}_{tst})}$ | RMSE |
| --- | --- | --- | --- | --- | --- | --- |
| Ad. Shr. | 102 | -581716.3 | 59.41 | 0.438 | 0.322 | 1.451 |
| Point Nor. | 106 | -583809.9 | 59.36 | 0.429 | 0.514 | 0.978 |
| Laplace | 100 | -584317.2 | 58.82 | 0.434 | 0.520 | 0.970 |

### Factor analysis identifies sets of compounds coordinated by biological processes

One possibility is that unobserved latent factors represent an underlying biological process that creates covariation among metabolites. Another possibility is that covariance among metabolites caused by population structure is captured by latent factors. We sought to partition latent factors into those due to a biological process and those due to a confounding effect (Bello et al. 2018). Since we showed that most population structuring of metabolites was caused by drift, we expect their coordination to be largely random, and therefore unrelated to their functional class. We assessed enrichment for functional classes within each factor, as well as the relationship between factors and population structure.

To assess biological enrichment, we determined whether the variance explained by a given metabolite functional class within a factor was significantly greater than might be expected by chance. The ontologies described in the preceding section were used to calculate the percentage of variance explained (PVE) by each functional class for each factor. To compute *p*-values, we compared these values against an empirical null distribution that was generated by randomly sampling loadings for a number of compounds equal to the number of compounds belonging to the functional class. This accounts for both the size of the class and the amount of variation that is explained by each factor. Of the 100 factors identified with the EBMF approach, 37 showed significant enrichment in one or more categories at the super-class level, while 40 and 36 factors showed significant enrichment at the class level and subclass levels, respectively (*q* < 0.05). Functional classes associated with lipids were most frequently enriched in our dataset (Fig. 2A,B), indicating that many factors may be capturing components of lipid metabolism. In addition to lipids, four factors showed enrichment for carbohydrates and carbohydrate conjugates, as well as amino acids. These results suggest that many latent factors are capturing meaningful biological processes that shape the seed metabolome, and can help shed light on the meaning of unannotated metabolites.

**Figure 2.**
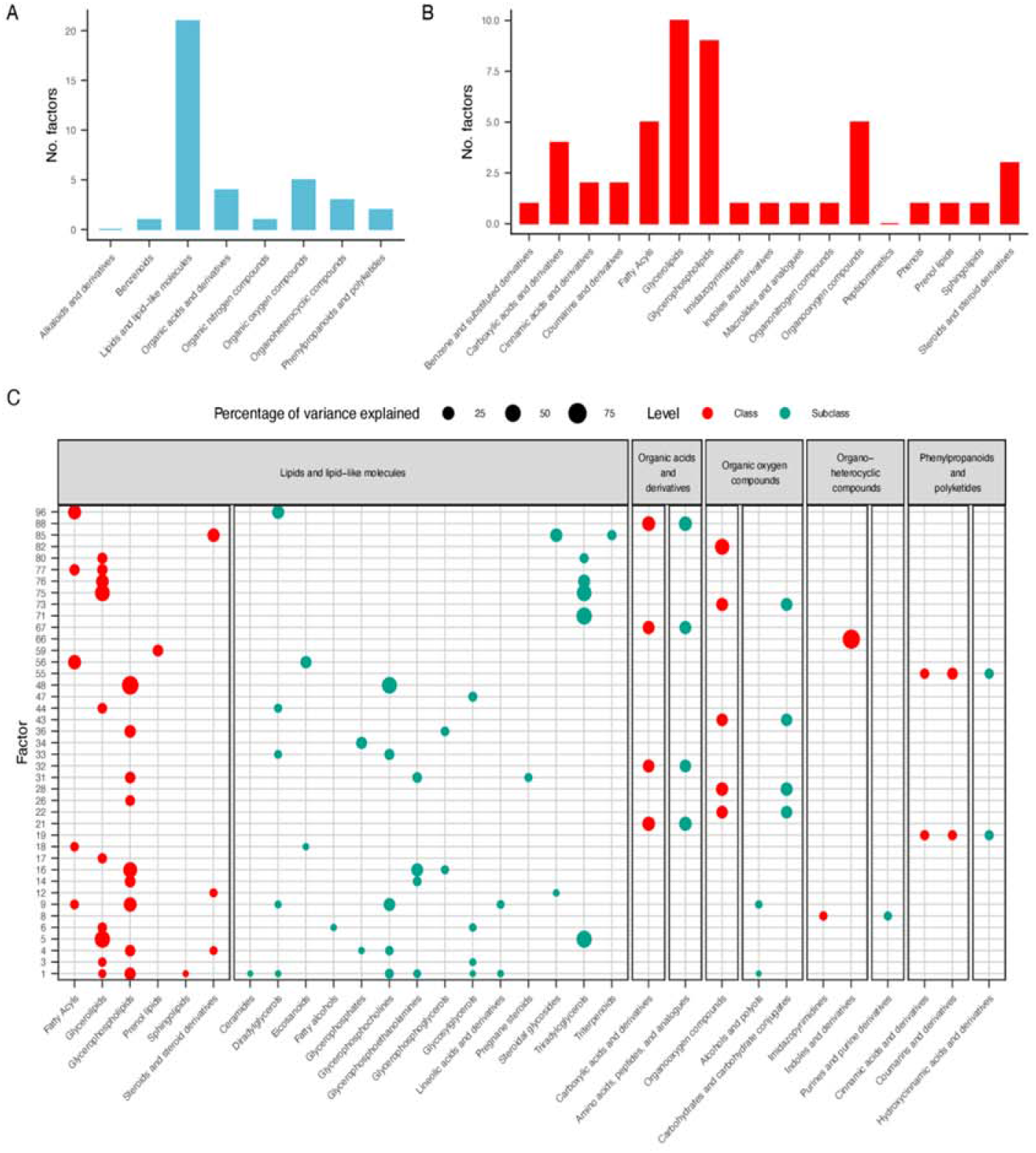
Functional enrichment among latent factors. Number of latent factors enriched (FDR < 0.05) for functional categories at the super-class level (A) and class level (B). Percentage of variance explained for each factor by a given functional category (C). Each point represents a functional class that was significantly enriched for one or more factors with the size of the point being proportional to the percentage of variance explained by that class for a given factor. Only factors and classes that showed significant enrichment (*q* < 0.05) at the super-class level are pictured. Colors differentiate between the class and subclass levels of the taxonomic hierarchy.

To address the possibility that latent factors are due to population structure, we examined the PVE by subpopulation. A linear model was fitted to each latent factor that included subpopulation assignment as a fixed effect. The PVE by subpopulation ranged from 0.03 to 29.8%, and subpopulation explained more than 20% of the variation for factors 7 and 12. Factor 7 did not show functional class enrichment but factor 12 was enriched across all hierarchies for lipid and lipid-like molecules — specifically steroidal glycosides (*q* < 0.05). Interestingly, *P*_*st*_ for this factor (0.27) was higher than the top 80th percentile of *F*_*st*_ (0.23), suggesting that the differences between subpopulations for this factor may be due to selection rather than drift. The high frequency of enrichment for functional classes of metabolites, and the relatively small amount of variation that was attributed to subpopulations suggests that these constructs can provide biochemically meaningful insights into the seed metabolome.

### Elucidating the origin of latent factors

To determine whether the covariance generated by each factor was due to genetic or environmental causes, we partitioned variance in latent factors into additive genetic and non-genetic components, and examined their genetic architecture. A Bayesian whole genome regression approach, Bayes C*π*, was used to estimate variance components, and estimate the degree of polygenicity of each factor (Habier et al. 2011). Bayes C*π* assumes markers have a zero effect with probability *π* and a non-zero effect with probability (1 − *π*). *π* is treated as an unknown and is estimated from the data. Thus, the magnitude of (1 − *π*) can provide a metric to assess the polygenicity of the trait. Narrow-sense heritability estimates (*h*^2^) ranged from 0.01 to 0.80, indicating that variation for many of the latent factors could be attributed to additive genetic effects (Figs S3,4). Moreover, the range of (1 − *π*) indicates this genetic variance is manifested in a wide range of architectures (Fig. S4).

The distribution of loading values for each latent factor was not similar — meaning that some factors show dense loadings (i.e., they generate covariance for many metabolites), while others show sparse loadings. These loadings are sampled from a scale mixture distribution where non-zero loadings are sampled from a Laplace distribution with a probability of (1 − *v*) and a point-mass at zero with a probability of *v*. Given that latent factors with dense loadings will generate covariance for many metabolites, we hypothesized that these factors will likely have a complex genetic architecture. To test this, we performed a partial Spearman’s correlation between polygenicity and the density of factor loadings while accounting for the heritability (*h*^2^) of each factor. A significant positive correlation between (1 − *v*) and (1 − *π*) was observed (ρ = 0.35; *p* = 4.5 × 10^−4^), indicating that factors that capture (co)variance among many metabolites tend to be controlled by many loci with small effects (Fig. 3; Figs S3,4). Several exceptions to this relationship were observed. For instance, factors 4, 13, 17, and 25 exhibited low polygenicity and dense loading patterns (Table II), indicating that these factors may be driven by loci with pleiotropic effects on the metabolome.

**Table II.**
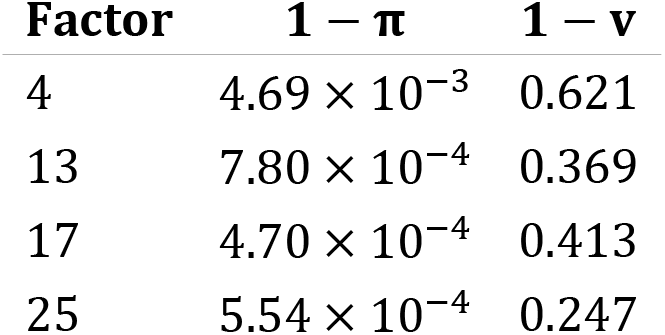
Factors capturing covariance between many metabolites with simple genetic architectures. Polygenicity estimates were based on the posterior means of 1-*π* and density of factor loadings are provided as 1 − *v*.

| Factor | $1 - \pi$ | $1 - v$ |
| --- | --- | --- |
| 4 | $4.69 \times 10^{-3}$ | 0.621 |
| 13 | $7.80 \times 10^{-4}$ | 0.369 |
| 17 | $4.70 \times 10^{-4}$ | 0.413 |
| 25 | $5.54 \times 10^{-4}$ | 0.247 |

**Figure 3.**
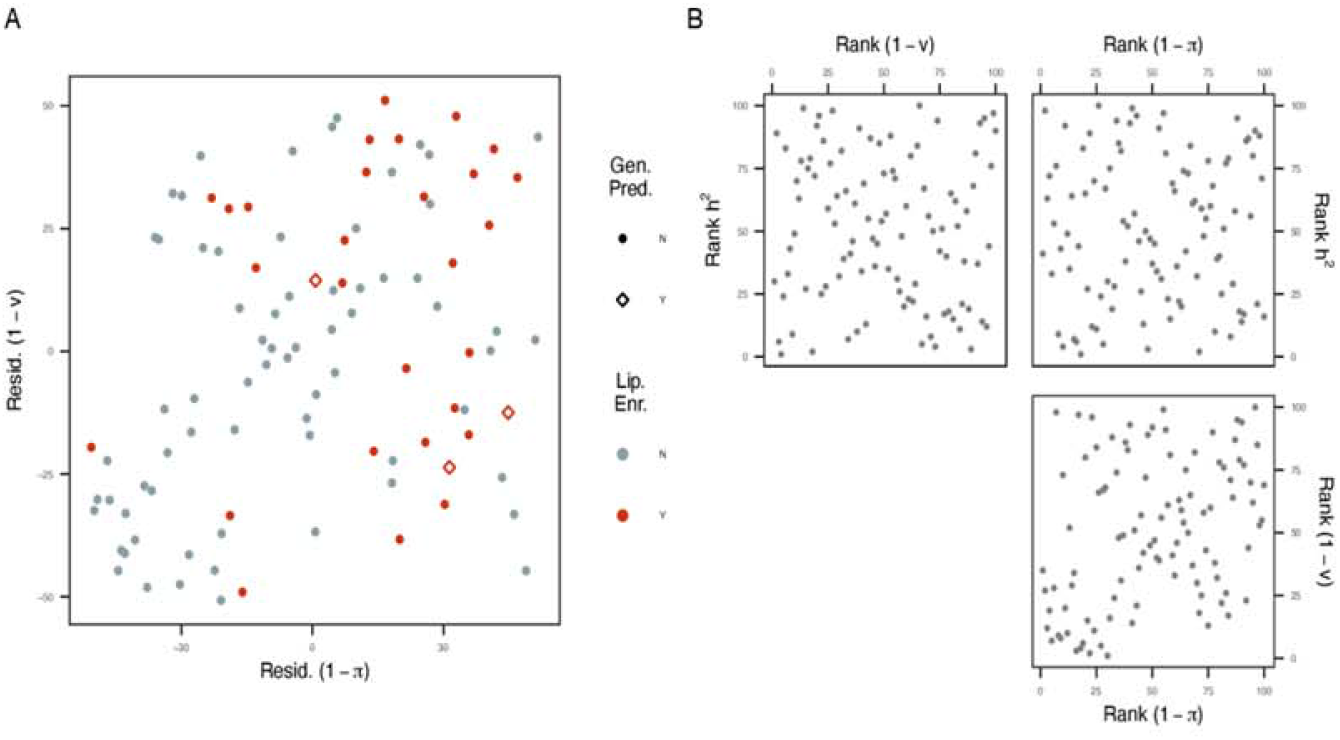
Relationships between polygenicity, density and heritability. (A) Association between polygenicity (1 − *π*) and density ranks (1 − *v*) after accounting for heritability (*h*^2^). Each variable was ranked from smallest to largest and the ranks for (1 − *π*) and (1 − *v*) were each regressed on ranks for *h*^2^. The scatter plot depicts the relationship between the residuals (Resid.) for each of these models. Colored points indicate factors that were enriched for lipids (Lip. Enr.), and different shapes indicate whether the factor was used to inform the lipid-enriched kernel for genomic prediction (Gen. Pred.). (B) Pairwise relationships between the ranks for each variable.

### Biologically-informed prediction of seed quality traits

Ultimately, the aim of this study was to translate insights from the metabolome into genetic resources that can be used by breeders to make broad changes to oat seed composition. In this respect, we assume that loci with large effects on multiple metabolites will be a more valuable resource to oat breeders than loci that affect one or a few metabolites. A conventional mixed linear model GWAS approach was used to identify loci with large effects on the latent factors. We identified 666 markers associated with 23 factors (*p* < 2.57 × 10^−7^; File S3; Figs S5-27).

We sought to address whether these associations could be leveraged to improve genomic prediction for seed quality traits in two independent studies. The first study quantified ten fatty acids (FA) in mature seed for 338 oat lines grown in two locations using targeted GC-MS. Of the 338 accessions evaluated, 330 overlapped with the factor analysis panel. The second study assayed seed lipid and protein content using near-infrared spectroscopy (NIRS) for 210 accessions from six trials with 12 lines overlapping with the factor analysis panel. Two prediction frameworks, genomic BLUP (gBLUP) and a multi-kernel prediction model (MK-BLUP), were used to predict seed-quality phenotypes across trials. The MK-BLUP framework uses two kernels to capture additive genetic effects, one of which is constructed from markers associated with latent factors and is referred to as the “biologically-informed" kernel. The second kernel is constructed from all other markers. Two biologically-informed kernels were evaluated: one that used markers associated with any latent factor to improve prediction, and one that only used markers associated with factors enriched for “Lipid and lipid-like molecules" (factors 4, 17, and 34). Prediction accuracy was assessed using five-fold cross validation with 50 resampling runs, and the MK-BLUP models were deemed to significantly improve prediction if prediction accuracies for MK-BLUP were higher than gBLUP in 90% of resampling runs.

Prediction accuracies were similar between gBLUP and the MK-BLUP models that used associations for all factors for nearly all traits. The exception was 18:3, which exhibited a 2.27% increase on average over gBLUP. The MK-BLUP approach significantly outperformed gBLUP for eight of the 12 traits considered when the kernel was informed by markers associated with lipid-enriched factors. For FA traits, the percent change in prediction accuracy over gBLUP ranged from −0.57% to 23.10%, with seven compounds showing significantly greater prediction accuracy compared to gBLUP (Fig. 4). Improvements were most notable for 14:0 and 16:0, which exhibited more than a 20% improvement over gBLUP. For NIRS traits, the lipid-enriched MK approach significantly improved predictions for lipid content on average by 9.9% (Fig. 5). These results show the latent factors and the genetic signals associated with them are reproducible and can be extended to new metabolite traits. Most importantly, these genetic signals are robust across populations and phenotyping technologies.

**Figure 4.**
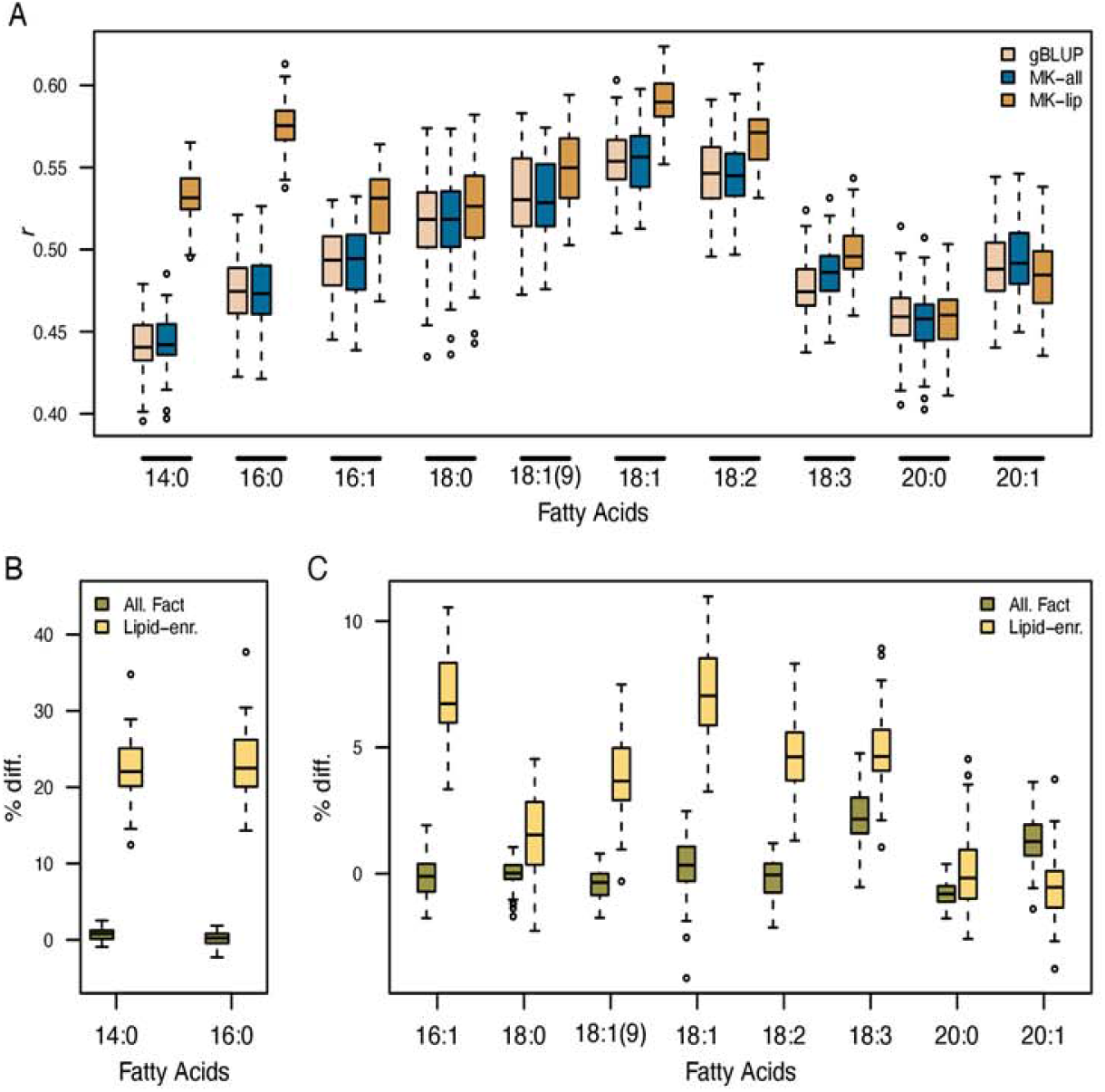
Genomic prediction for fatty acid compounds. Prediction accuracy was assessed using five-fold cross validation with 50 resampling runs. (A) The distribution of Pearson’s correlation (*r*) coefficients between observed phenotypes and genetic values for each fatty acid compound. Panels B and C show the percent difference (% diff.) in prediction accuracy for the multikernel (MK) approach relative to genomic BLUP (gBLUP). The suffixes ‘-all’ and ‘-lip’ indicate models where the biologically-informed kernel was constructed from markers associated with any latent factor or lipid-enriched factors, respectively.

**Figure 5.**
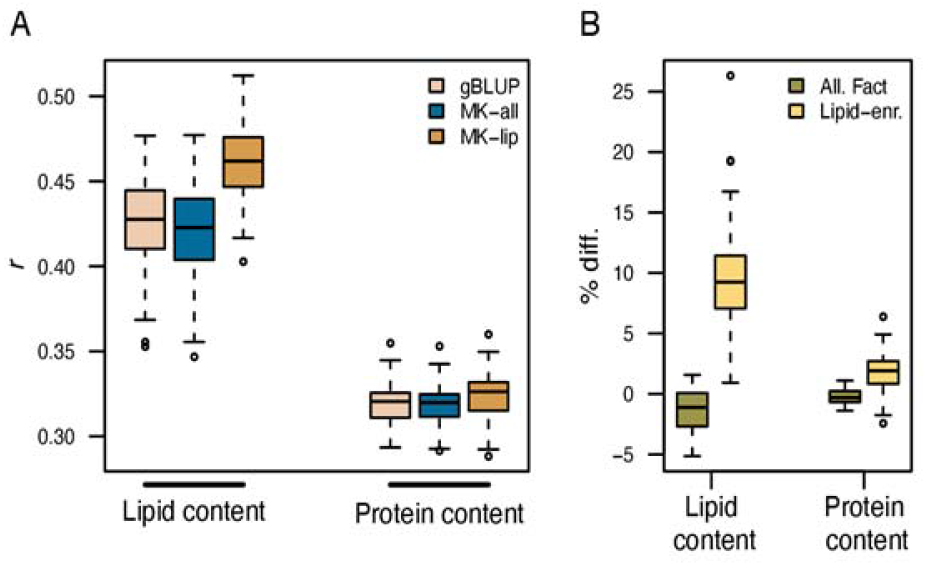
Genomic prediction for lipid and protein content measured via NIRS. Prediction accuracy (*r*) was assessed using five-fold cross validation with 50 resampling runs. Panel A shows the distribution of Pearson’s correlation coefficients between observed phenotypes and genetic values. Panel B shows the percent difference (% diff.) in prediction accuracy for the multikernel (MK) approach relative to genomic BLUP (gBLUP). The suffixes ‘-all’ and ‘-lip’ indicate models where the biologically-informed kernel was constructed from markers associated with any latent factor or lipid-enriched factors, respectively.

## Discussion

The oat seed harbors a rich array of biochemical compounds that are important for human health, and considerable variation for these compounds exist in oat germplasm (Peterson and Wood 1997; Frey and Holland 1999; Gulvady et al. 2013; Zhou et al. 2019). Efficiently accessing this variation is necessary to rapidly deliver oat varieties with beneficial nutritional profiles to the consumer. Advances in metabolic profiling over the past 20 years has provided a suite of tools to comprehensively assess these compounds, along with many others, in large populations and to elucidate their regulation (Keurentjes et al. 2006; Tohge and Fernie 2010). Structural elucidation and metabolite identification remain a significant bottleneck in characterizing the metabolome using untargeted metabolomics (Dunn et al. 2013). Many of the publicly available databases do not adequately capture the rich diversity of metabolites that are produced in plant species (De Vos et al. 2007; Tohge and Fernie 2010). Therefore, approaches that uncover the relationships between metabolites, both known and unknown, may help shed light on the function of these compounds.

Despite being able to reliably detect the abundance of 1,668 compounds in the current study, less than a third of these compounds were annotated. We used a latent factor approach that leverages the correlation between metabolites to help elucidate their function. Our rationale is that metabolites that participate in the same pathway should be correlated. Thus, by extracting the major correlation patterns in the observed variables we can begin to elucidate the biochemical pathways that shape the seed metabolome. Moreover, by studying the relationships among annotated metabolites, we can generate new hypotheses to understand the function of unannotated compounds.

### Characterizing the metabolome using latent factors

Our enrichment approach helped shed light on the biochemical processes these latent factors might affect. For instance, we found significant enrichment for a range of processes associated with primary metabolism (amino acids, phospholipid metabolism) and secondary metabolism (coumarin and terpenoid metabolism). Since roughly 30% of the metabolites assayed had functional annotations, this enrichment approach may shed light on the function of unannotated metabolites. For instance, factor 4 showed significant enrichment for “lipid and lipid-like molecules.” Although, only 45 of the top 100 compounds with high loadings were annotated, the high correlation between these unknown compounds and annotated, lipid-like compounds suggests putative role in lipid metabolism. Although further analyses are necessary to elucidate the structure of these unknown metabolites, our enrichment approach provides a data-driven approach to generate hypotheses for these unannotated metabolites.

One overarching pattern observed across latent factors is the enrichment for compounds related to lipid metabolism. At the most coarse level, super-class, 21% of factors were enriched for “lipid and lipid-like molecules,” and these patterns were consistent at more specialized levels of lipid metabolism. Oat is unique among cereals in both the abundance and distribution of lipids within the seed (Price and Parsons 1975; Gulvady et al. 2013; Frey and Holland 1999). And with approximately 57% of the annotated metabolites in our data classified as lipid-like compounds, it is not surprising that categories associated with lipid metabolism were most frequently enriched.

It is possible that other processes are prevalent in the metabolome and are reflected in the latent constructs, but remain undetected due to the annotations that were used for functional enrichment. These ontologies are based on structural similarities between compounds rather than pathway-based relationships. We expect compounds involved in the same pathway to be correlated, and since latent factors are defined by these correlations they should in some sense be an abstraction of these pathways. Biochemical reactions often involve compounds with dissimilar structures, thus enrichment based on structural similarities may bias enrichment towards pathways composed of structurally similar metabolites (e.g., lipid metabolism). While this enrichment approach may be imperfect, other studies have used similar approaches and have proven to be useful in other species (Barupal and Fiehn 2017; Fan et al. 2018; Marco-Ramell et al. 2018; Showalter et al. 2019). The ChemRich approach developed by Barupal and Fiehn (2017) uses the ClassyFire ontology to classify compounds into functional classes and tests for enrichment using a Kolmogorov–Smirnov test. Annotations that map metabolites to a pathway can provide additional evidence that these latent factors are indeed due to an underlying biochemical process; however, current resources do not provide the breadth and resolution necessary to perform such analyses.

### Understanding the origin of latent factors

Although it may seem reasonable to suggest that the observed covariance among metabolites is due to a biological cause that is manifested in the metabolome, making causal inferences from observational data is nontrivial due to the presence of confounding factors (Spirtes et al. 2000; Rosa and Valente 2013; Bello et al. 2018). Given these data were collected on a structured population, it is expected that some of this covariance can be attributed to population structure. This can influence the construction of latent variables (Phillips et al. 2001) if not taken into account. There are many ways to account for structure in the definition of latent factors, either by including the genomic relationship matrix, or some component(s) of it, in the factor analytic model or by regressing-out these effects prior to factor analysis; however, it is important to consider whether these steps are necessary. While such measures will control for confounding due to structure, they will also remove possibly meaningful biochemical relationships that are associated with structure. If a set of compounds participating in a common pathway happen to differ between subpopulations, correcting for structure may remove the latent factor that describes this process. We identified two latent factors, factors 7 and 12, that were highly associated with population structure. Enrichment analysis and *P*_*st*_ − *F*_*st*_ suggested that factor 12 may indeed describe a biological process (steroidal glycoside metabolism) that was affected by selection. This factor would likely be removed if structure were accounted for prior to factor analysis.

If subsequent genetic analysis are planned for latent factors, regressing-out structure may also remove meaningful genetic signal. Given the minor structure observed among accessions in the diversity panel and the importance of preserving genetic signal in the factor scores, we thought that measures to account for structure could be harmful to the study as a whole. Moreover, our downstream association mapping approaches accounted for population structure by using the first two PCs and a kinship matrix based on allele dosages. In the event that some latent factors were defined based on kinship, we do not expect to recover any signal from association mapping with scores for these latent factors.

We should not place too much emphasis on causality in a purely biological sense when interpreting these latent factors. Rather it is important to consider the limitations of the study, interpret latent factors with caution, and view them as a means to generate testable hypotheses. The aims of our study were to (1) elucidate the major biochemical processes in the oat seed metabolome, and (2) to leverage these insights to improve selection for seed quality. Thus, hypotheses are generated in the former and are tested in the latter. If latent factors do not represent a causal effect then we should not see any improvement in predictions when inferences on these constructs are extended to new studies and/or populations.

### Translating ‘omics’ insights to crop improvement

Two independent studies were used to determine whether biological signal in the latent factors could be generalized to other populations and/or traits. The fatty acid dataset can be used to test whether the information learned by latent factors is reproducible, while the NIRS dataset provides a means to test whether this information is transmissible to related traits in new populations. We distinguish between these two because: (1) the majority of accessions included in the fatty acid dataset are accessions that were used for the factor analysis metabolome study, while less than 6% of accessions are common between the factor analysis and the NIRS studies; (2) the fatty acid data was generated using targeted metabolomics, meaning there should be a high correspondence between the metabolites measured in the fatty acids study and those that were assayed for the factor analysis metabolome study (Carlson et al. 2019).

Considering these aspects, we expect that the information learned from the factor analysis metabolome study should have the most pronounced effect on predictions for fatty acid compounds. Consistent with these expectations, we observed the greatest improvements in prediction accuracy among all traits for the biologically-informed prediction model over gBLUP for these compounds when the kernel was constructed using associations for lipid-enriched factors. Thus, the genetic signal that is associated with these latent factors is relevant to both studies and phenotyping approaches (i.e., targeted and untargeted metabolomics). A comparison of the GWAS hits in (Carlson et al. 2019) and those in our study showed little overlap, with two common associations identified for factor 13 and the tenth PC of fatty acid phenotypes in (Carlson et al. 2019), and factor 17 and 14:0 in (Carlson et al. 2019). Of these two factors, only factor 17 showed enrichment for “lipid and lipid like molecules” at only the super-class level. While *q* values at more specific functional classes were above the chosen significance threshold, *q* < 0.05, enrichment for 1-acyl-sn-glycero-3-phosphocholines was the top-ranked category at the parental class (*q* = 0.058). Interestingly, hydrolyzation of these compounds by phospholipase A1 yields a fatty acid. Although additional studies are necessary to elucidate the biochemical pathways associated with factor 17, these results provide an interesting link between 1-acyl-sn-glycero-3-phosphocholines catabolism and fatty acid abundances and the possibility of modifying 1-acyl-sn-glycero-3-phosphocholine metabolism to fine-tune fatty acid profiles in oat. Although it is difficult to connect loci associated with latent factors with changes in specific metabolites, our polygenicity analysis offers a more general explanation – specifically, that these loci may affect many metabolites.

The second study with NIRS-derived composition measurements provides several realistic challenges, and should be a reasonable estimate of how the biologically-informed model would perform in a breeding program. The population that was evaluated for NIRS phenotypes is largely independent from the population that was used for factor analysis. Moreover, the NIRS phenotypes are only approximations of total lipid or protein content. The advantage of using NIRS to estimate seed metabolites is that it is a relatively low cost phenotyping approach compared to metabolomics and is high-throughput, making it a tractable solution for many breeding programs interested in improving health-promoting compounds (Diepenbrock and Gore 2015). Despite these challenges the multi-kernel prediction approach – when informed using markers associated with lipid-enriched factors – significantly improved prediction for lipid content compared to gBLUP.

### On the relationship between factor density and polygenicity

The positive relationship observed between the magnitude of polygenicity and loading densities, indicates that latent factors that influence many metabolites are more likely to have a complex genetic architecture. These observations are somewhat expected. If these dense latent factors are representative of some central component of the metabolome, perturbations on these processes would likely result in large-scale biochemical changes that may affect fitness. Therefore, it is important that these processes are robust to mutations and are maintained at, or near some optima. This is the basis of canalization – important physiological processes will evolve to reach robust optima – and suggests that much of the oat seed metabolome is under optimizing or stabilizing selection (Gibson 2009; Slatkin 1970; Waddington 1942).

Perhaps what is more interesting is the factors that deviate from this relationship, specifically factors 4 and 17. Both exhibited dense loading patterns, oligogenic architectures (ranked 8th and 17th for density, respectively, and 50th and 73rd for polygenicity), and were enriched for lipids. The large-effect loci associated with these latent factors may have pleiotropic effects, or may consist of a set of tightly linked genes that influence the abundance of lipid-like compounds. In either case, this may explain the deviance from the density-polygenicity relationship observed for other factors. The presence of these loci raises a larger question, specifically *Why are these loci segregating in the population?* The theoretical and simulation studies by Orr, as well as empirical evidence in maize and other species may help explain these observations (Orr 1998; Orr 1999; Boyko et al. 2010; Brown et al. 2011; Carlborg et al. 2006; Colosimo et al. 2004; Doebley et al. 1997; Van Laere et al. 2003; Wang et al. 2005). For “older” traits – *i.e.* those associated with adaptation in natural environments – such large effect alleles at these loci would likely be removed through negative selection as these alleles may shift phenotypes far from the optimal values (Orr 1998; Orr 1999). This was proposed by Brown et al. (2011) to explain the small effect sizes for flowering and leaf traits in maize. This is not necessarily the case for traits that are relatively “new” in evolutionary history or are not associated with adaptation. For instance, plant architecture and inflorescence traits have relatively simple genetic architectures in maize and are recent targets for artificial selection (Brown et al. 2011; Doebley et al. 1995; Doebley et al. 1997; Wang et al. 2005; Wallace et al. 2014). This is also the case for traits under recent artificial selection in other species (Boyko et al. 2010; Carlborg et al. 2006; Colosimo et al. 2004; Van Laere et al. 2003). While it is unknown whether seed lipid content has any adaptive significance in oat, lipid content and traits that are genetically correlated with lipid content (i.e., β-glucans) are popular targets for many breeding programs (Welch and Lloyd 1989; Kibite and Edney 1998; Cervantes-Martinez et al. 2002). Thus, the oligogenic architectures for factors enriched for lipids may be a reflection of this relatively recent selection by breeders for lipids or traits that are genetically correlated with lipids.

## Conclusions

This study shows that we can translate biological knowledge obtained from the characterization of high dimensional ‘omics’ data to improve prediction and selection for agriculturally important traits. The matrix factorization approach used here provides an effective means to reduce the dimensionality of the data while preserving important biological features that generate correlation in the observed phenotypes. This can help reduce the multiple testing burden often experienced with GWAS on ‘omics’ data and allow the recovery of meaningful genetic signal. This signal can be leveraged to improve prediction for low-cost phenotypes that provide an approximation of biochemical attributes in independent populations. In a broader context, this approach that can be used to manage the allocation of phenotyping resources and improve breeding decisions. For instance, breeders can phenotype a single replicate of a ‘discovery’ population with a costly, high-resolution ‘omics’ technology and these data can be used to inform predictions for low-cost, lower-resolution phenotypes in new populations or trials. These approaches can be easily extended to other crops, tissues and ‘omics’ technologies to improve predictions for complex traits.

## Materials and Methods

### Plant materials and growth conditions

The oat diversity panel consists of 375 accessions derived from breeding programs in North America and Europe. In 2018, the panel was grown in an augmented field design in Ithaca, NY, which consisted of 368 unreplicated entries allocated to 18 blocks with 21-23 plots per block. One primary check, ‘Corral’, was included in each of the blocks, while one of six secondary checks were randomly allocated to each block. These secondary checks were replicated four times, while the primary check was replicated 19 times (one block had two ‘Corral’ plots).

### Latent factor analysis

Latent factor analysis seeks to identify a set of *k* unobserved, latent factors that give rise to the observed covariance among a set *p* of observed variables. Formally, this relationship is given by

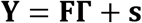

where **Y** is a centered and standardized *n* × *p* matrix of observations for *p* metabolites and *n* individuals; **F** is an *n* × *k* matrix of factor scores; **Γ** is a *k* × *p* matrix of loadings; and **s** is an *n* × *p* matrix of specific effects. The (co)variance matrix **V** of observations **Y** is decomposed into common covariance and specific covariance:

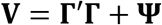

All matrices are defined as above, and **Ψ** is a *p* × *p* diagonal matrix of specific variances.

A recent framework described by Wang and Stephens (2018) uses an empirical Bayes framework to learn appropriate priors from the data given a family of densities. This approach, Empirical Bayes Matrix Factorization (EBMF), can tailor the sparsity for factor loadings and scores based on what best fits the data. This was implemented using the `flashr` package in R (https://github.com/stephenslab/flashr). Three classes of models were fit that differed in families of densities used to fit the data: Laplace, point-normal, and adaptive-shrinkage. A combination of the ‘Greedy’ search algorithm and backfitting was used to define the model.

We evaluated the classes of models for goodness-of-fit using percent variance explained (PVE) by the common factors and predictive ability using three-fold orthogonal cross validation (3-OCV) (Owen, et al. 2016). PVE was defined as

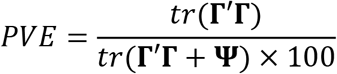

with *tr* indicating trace of the given matrix and all other matrices defined as above. 3-OCV is similar to classical CV, but ensures that no rows and columns of the testing data (**Y**_**test**_) have all missing data. The model above was fitted for the training set data and predicted values for the testing set were calculated via 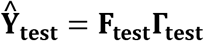. The accuracy of each model was evaluated using the root mean square error (RMSE) and the correlation between predicted and observed values for observations in the testing set for each fold. Ten independent resamplings were performed. The metrics were averaged over folds, and the ‘best’ model was selected based on the results across the ten repeats.

### Enrichment analysis for latent factors

We used the ClassyFire taxonomic hierarchies to test for functional enrichment for each factor (Feunang et al. 2016). Briefly, ClassyFire uses a hierarchy of five levels to describe chemical compounds. At each level, we calculated the percentage of variance explained (*PVE*_*kc*_) for factor *k* by functional class *c*. This is given below

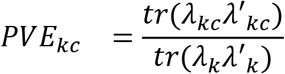

where *λ*_*k*_ is a vector of loadings for a given factor *k*, and *λ*_*kc*_ is a vector of loadings of factor *k* for compounds in class *c*. Our null hypothesis is that the variance captured by compounds in a given class will be equivalent to that explained by a random set of compounds of equal size to that class. To test this, we generated an empirical null distribution for each functional class and factor. For each class and factor, we picked a random set of compounds with a size equivalent to the class by sampling the loadings of 1,668 metabolites without replacement and computed PVE. This process was repeated 1,000 times for each combination of functional class and factor. For each class-factor combination, we compared observed PVE with the empirical null distribution for that given combination and calculated *p*-values. Finally, to account for multiple testing, *q*-values were calculated across all factors and classes following (Storey 2002). Functional classes with fewer than five compounds were excluded from analyses to ensure that results were not biased to small classes with one or two compounds with very high loadings.

### Assessing the genetic architecture of latent factors

#### Genome-wide association study

To identify loci associated with latent factors, the following linear mixed model was fit to factor scores for each latent factor (*k*)

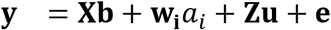

where **y** is a vector of factor scores; **X** is a matrix of the first two PCs and **b** is the corresponding vector of effects; **w**_**i**_ is a vector of allele dosages for marker *i* and *a*_*i*_ is the corresponding marker effect; and **u** is a vector of polygenic effects. The first two PCs explained about 13% of the genomic relatedness among lines. We assume 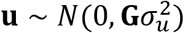 and 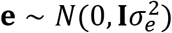, where **G** is a genomic relationship matrix calculated following the second definition provided by VanRaden (2008). These models were fitted using the `rrBLUP` package in R (Endelman 2011). GWAS was performed using 62,049 SNP markers with a minor allele frequency > 0.05 and 335 individuals with marker data and factor scores.

We used the approach described by J. Li and Ji (2005) to account for multiple tests performed both within and across factors. Briefly, we computed the number of effective tests (*M*_*eff*_) by performing eigenvalue decomposition on the correlation matrix for 62,049 markers. This provides an estimate of the number of tests performed within each factor.

Next, we multiplied this value by the total number of factors. The test criteria was then adjusted using *M*_*eff*_ with the Sidak correction below (Šidák 1967).

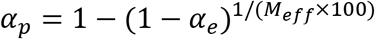

This provided a genome-wide significance (*α*_*p*_) value of 2.57 × 10^−7^ at *α*_*e*_ = 0.1 with *M*_*eff*_ = 4,097.

#### Estimating polygenicity with Bayes Cπ

To estimate polygenicity of each factor, we used Bayes C*π* (Habier et al. 2011). Bayes C*π* is a Bayesian whole-genome regression approach that can be used to estimate the proportion of markers with a non-zero effect on the phenotype. Bayes C*π* assumes that marker effects are drawn from a mixture distribution. Effects drawn from a distribution with a point mass at 0 with a probability *π* and a univariate Gaussian distribution with probability (1 − *π*). The linear model is given below.

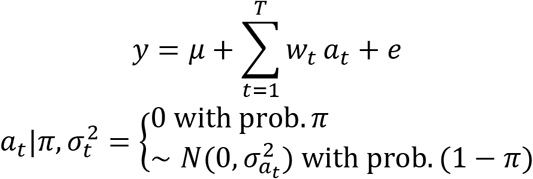

*w*_*t*_ is a vector of marker genotypes for marker *t* and *a*_*t*_ is the corresponding effect. The above model was fitted using the `JWAS` package in Julia using factor scores and 62,049 markers (Cheng et al. 2018). We used 200,000 iterations and discarded the first 100,000. Posterior means of 1 − *π* were used as estimates of polygenicity.

### Genomic prediction of seed quality traits

Two studies were used to determine whether associations from factor score-based GWAS could improve genomic prediction accuracies. The first consisted of fatty acid measurements for 500 lines, of which 338 had corresponding genotypic data consisting of 61,900 markers. These lines were evaluated at two locations in New York in 2014 (Carlson et al. 2019). The second consisted of six trials that evaluated protein and lipid content using near-infrared spectroscopy for 210 lines, of which 12 overlapped with the lines used for factor analysis. For this study 58,293 markers were used for prediction. Table S2 lists the trials used for genomic prediction and links to access these data.

A multi-kernel BLUP model was used to predict seed phenotypes across trials. Additive genetic effects were predicted using two kernels. The first is computed using markers that were identified through factor score-based GWAS and is referred to as the biologically-informed kernel, while the second was computed using all other markers. This model is given by

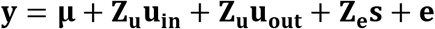

where **y** is a vector of phenotypes; **Z**_**u**_ is an *n* × *q* incidence matrix that assigns the *q* genomic values to *n* observations; **u**_**in**_ and **u**_**out**_ are genomic values predicted from biologically-informed or non-informed kernels, respectively; and **Z**_**e**_ is an *n* × *e* incidence matrix that assigns observations to trials and *s* are the corresponding effects. Moreover, we assume 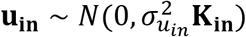, 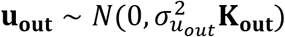, and 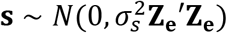. Where **K**_**in**_ and **K**_**out**_ are biologically-informed and non-informed kernels genomic relationship matrices, respectively, and are computed according to VanRaden (2008). We considered two marker sets to compute these matrices: markers associated with any latent factor, and markers that were associated with latent factors showing enrichment for lipid and lipid-like molecules at the superclass level (*q* < 0.05). Markers that were in weak linkage disequilibrium (LD) (*r*^2^ > 0.25) with GWAS hits were included in the biologically-informed kernel. LD was computed separately for each study.

Genomic BLUP (gBLUP) served as a base-line to compare the ability of the multi-kernel approach to predict seed phenotypes. The model is similar to the multi-kernel model; however, the relationship matrix was constructed using all available markers for each study. All models were fit using the `BGLR` package in R, with 20,000 iterations, of which the first 5,000 were discarded (Perez and de los Campos 2014).

Prediction accuracy was assessed using five-fold cross validation with 50 resampling runs, and was computed using Pearson’s correlation between observed phenotypes and predicted genomic values for accessions in the testing set. Genomic values for the multi-kernel approach were computed as the sum of breeding values from each random genetic effect. Correlation coefficients were averaged across folds.

## Data availability

All metabolomic data are provided via Cyverse and can be accessed using the following url https://de.cyverse.org/de/?type=data\&folder=/iplant/home/mcampbell4. All R code used for analyses is provided as Rmarkdown files and can be accessed via https://github.com/malachycampbell/OatLatentFactor.

## Acknowledgements

Funding for this research was provided by USDA-NIFA-AFRI 2017-67007-26502. Mention of a trademark or proprietary product does not constitute a guarantee or warranty of the product by the USDA and does not imply its approval to the exclusion of other products that may also be suitable. The USDA is an equal opportunity provider and employer.

## Supplemental Data

- **Supplemental Methods**
- **Supplemental Table S1.** Genotyping-by-sequencing experiments in Triticeae Toolbox used in this study.
- **Supplemental Table S2.** Trials in Triticeae Toolbox used for genomic prediction.
- **Supplemental Figure S1.** Summary of subpopulation clusters based on major geographic regions.
- **Supplemental Figure S2.** Three-fold orthogonal cross validation results for three EBMF approaches.
- **Supplemental Figure S3.** Spearman correlation between factor density (1 − *v*), polygenicity (1 − *π*) and narrow sense heritability (*h*^2^).
- **Supplemental Figure S4.** Distribution of density (1 − *v*), polygenicity (1 − *π*) and narrow sense heritability (*h*^2^) estimates.
- **Supplemental Figures S5-27.** Manhattan plot for factor score-based GWAS.
- **Supplemental File S1.** Deregressed best linear unbiased predictions for 1,668 metabolites.
- **Supplemental File S2.** Metabolites showing significant differences between subpopulations.
- **Supplemental File S3.** GWAS summary statistics

